# Automated structure refinement of macromolecular assemblies from cryo-EM maps using Rosetta

**DOI:** 10.1101/050286

**Authors:** Ray Yu-Ruei Wang, Yifan Song, Benjamin A Barad, Yifan Cheng, James S Fraser, Frank DiMaio

## Abstract

Cryo-EM has revealed many challenging yet exciting macromolecular assemblies at near-atomic resolution (3-4.5Å), providing biological phenomena with molecular descriptions. However, at these resolutions accurately positioning individual atoms remains challenging and may be error-prone. Manually refining thousands of amino acids – typical in a macromolecular assembly – is tedious and time-consuming. We present an automated method that can improve the atomic details in models manually built in near-atomic-resolution cryo-EM maps. Applying the method to three systems recently solved by cryo-EM, we are able to improve model geometry while maintaining or improving the fit-to-density. Backbone placement errors are automatically detected and corrected, and the refinement shows a large radius of convergence. The results demonstrate the method is amenable to structures with symmetry, of very large size, and containing RNA as well as covalently bound ligands. The method should streamline the cryo-EM structure determination process, providing accurate and unbiased atomic structure interpretation of such maps.

## Introduction

Advances in direct electron detectors as well as better image analysis algorithms have led cryo-electron microscopy (cryo-EM) to achieve near-atomic resolution (3-4.5 Å) using single-particle analysis [1-3]. Cryo-EM reconstructions at these resolutions, where individual β-strands are resolvable, and bulky sidechains are somewhat visible, make it possible to build an all-atom model directly from such maps [4,5]. Although sequence can be registered, density maps at this range of resolution do not grant enough information to precisely assign coordinates for each atom in the structure, from which molecular interactions for a biochemical process is captured. Furthermore, such model building and refinement is challenging and error prone [6,7]. Determination of detailed atomic interactions from these sparse sources of data is desirable, however, the inherent ambiguity in the data makes identifying these interactions extremely difficult, even for experts.

Model-building into a cryo-EM map at near-atomic resolution generally involves manually building a model into the map using a graphical user interface tool [8] followed by refinement with software repurposed from X-ray crystallography [9,10]. This process requires identification of key amino acid sidechains to register stretches of sequence within the map (possibly aided by the topology from a homologous structure), followed by extension of these short fragments of sequence to form one or more fully connected protein chains. At near-atomic resolution, this manual model-building and refinement can be error prone owing to: a) the density may not be of sufficient resolution to uniquely identify sidechain rotamers, even for bulky aromatic residues, making it difficult to accurately determine sidechain-sidechain or sidechain-backbone interactions; b) for regions of non-regular secondary structure (turns or loops) or with poor local resolution, it may be difficult to accurately position backbone atoms; and c) in these same regions, precise sequence registration may also be error prone. Getting these atomic interactions correct is crucial for understanding detailed atomic mechanisms of proteins, designing drugs with a very specific shape complementarity, and for understanding subtle conformational changes of a protein. A structure refinement procedure that can automatically improve the atomic details of a model from such density data is thus very much desired.

In this manuscript, we develop a three-stage approach for automatically refining manually traced cryo-EM models (Figure 1). While previously we have developed an iterative local rebuilding tool capable of refining homology models into near-atomic-resolution cryo-EM maps [11], several advances were required for extending this tool to successfully refine hand-built models. Our new approach includes a method for automatically detecting and correcting problematic residues in hand-built models without overfitting, a model-selection method for identifying models with good agreement to the density data and with physically realistic geometry, a voxel-size refinement method for correcting errors in calibrating the magnification scaling factor of a microscope, a novel sidechain-optimization method to correct sidechain placement errors in very large systems, and a way to estimate uncertainty in a refined model. These methods, combined, allow to correct backbone errors that significantly deviate from the starting model, but may still assign a high degree of confidence to these regions in the refined model.

**Figure 1.**
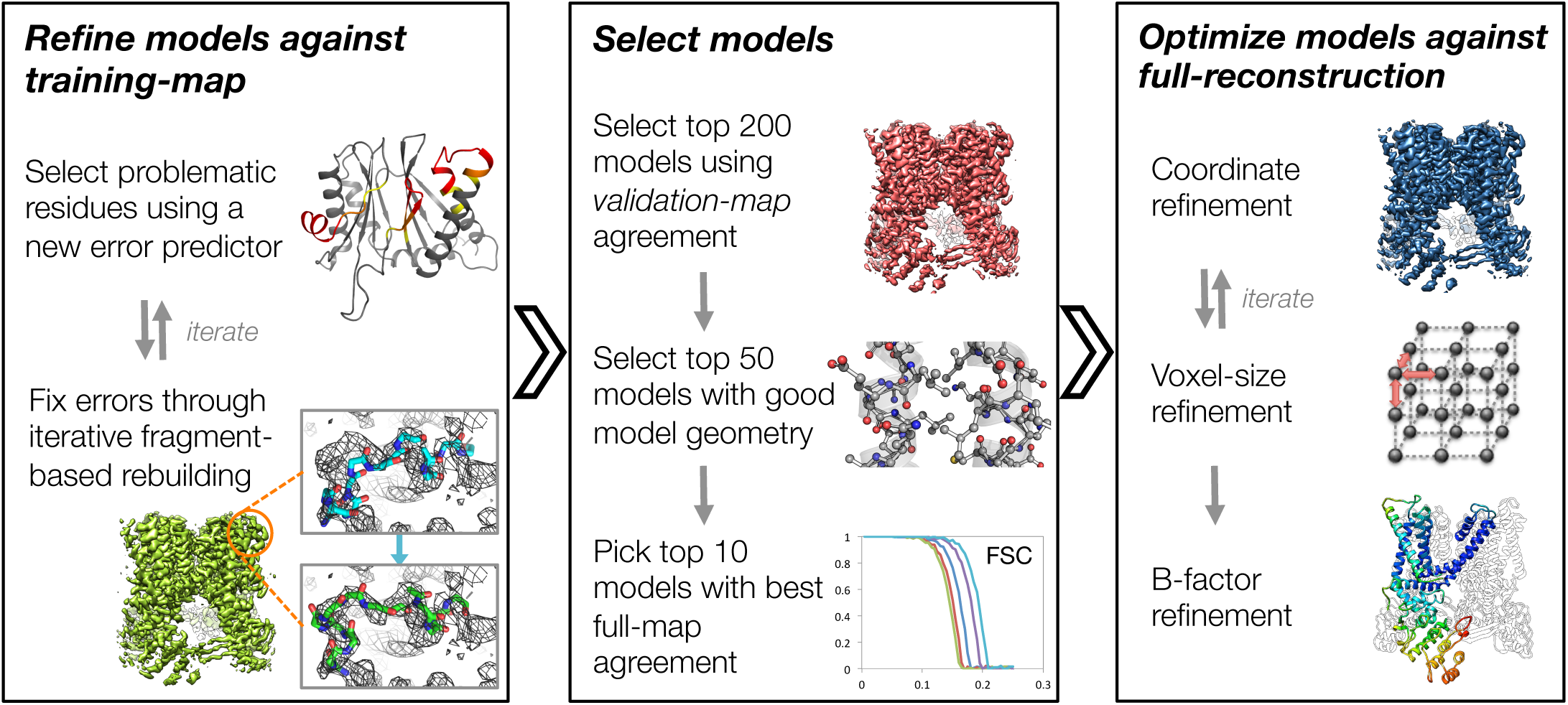
An overview of the three stages of automated refinement. (Left) In stage 1, problematic regions are predicted using a newly developed error predictor judging on local strain in the model and poor local density fit. These selected regions are subject to iterative fragment-based rebuilding with in a Monte Carlo sampling trajectory. Refinement in this stage is restricted to using one-half of the data, referred to as the training map. (Middle) In stage 2, the best models from the ~5000 independent Monte Carlo trajectories are selected. Models are selected based on: agreement to the validation map (independently constructed from the other half of the data), then by model geometry as assessed by MolProbity, and finally, based on agreement to the full reconstruction. At this point, the selected models should in general have good fit-to-density and good geometry without overfitting to the data. (Right) In stage 3, using the 10 best models selected, we then optimize against the full reconstruction. Two half maps are used for choosing the optimal density weight to refine structures using full-reconstruction. Finally, these top 10 models are optimized (without large scale backbone rebuilding) into the full-reconstruction, which alternates with voxel size refinement iteratively. Finally, these models are subject to B-factor refinement.

Finally, we apply this approach to three recently solved cryo-EM single particle reconstructions at near-atomic resolution: the TRPV1 channel at 3.4-Å resolution (TRPV1) [12], the F_420_-reducing [NiFe] hydrogenase (Frh) at 3.4-Å resolution [13], and the large subunit of mitochondrial ribosome at 3.4-Å resolution (mitoribosome) [14]. We show that in all three cases of diverse and large systems, we are able to automatically refine models to high-quality (as assessed by MolProbity), while maintaining or improving agreement to the density data. Significantly, in the case of TRPV1, we newly identify a biological relevant atomic interaction – a disulfide bond – not built in the originally deposited model, but supported in the literature. In the case of Frh, we show our refinement procedure led to 13.2% fit-to-density improvement mainly through optimization of the voxel size. Finally, in the case of mitoribosome, we show significant improvement in model geometry: the number of “Ramachandran favored” residues increases by 5%, and Molprobity [15] score improvement is observed in all 48 protein chains.

## Results

An overview of our refinement approach is shown schematically in Figure 1 (and is fully described in *Methods*). Broadly, the approach proceeds in three stages. In the first stage, we identify problematic residues by assessing local model-strain and local agreement to density data. These regions are rebuilt against a “training” half-map using fragment-based Monte Carlo sampling with many independent trajectories followed by all-atom refinement. Secondly, the best subset of these independent trajectories are selected by identifying a subset of stereochemically correct models with best agreement to an independent “validation” half-map, to prevent overfitting. Finally, models are further optimized in the full-reconstruction with a weight optimally scaled between experimental data and the forcefield using the “validation” half map. Our approach adopts and improves upon our previous work on refining cryo-EM structures from distant homology structures [11], in which a similar fragment-based backbone rebuilding strategy is employed. However, several critical improvements were necessary in extending our previous work to successfully refine hand-traced models, larger complexes, and a more diverse set of systems.

### Identification of backbone errors using local strain

In previous work [11], local fit to density is used to identify residues in a distant homology model to rebuilt. However, unlike remote homology models, hand-traced models typically fit the data very well, but are incorrect geometrically (strain). Consequently, a key improvement is to make use of model strain as a criterion in selecting regions to refine. Moreover, when the previous approach was applied to the *de novo* hand-traced models from cryo-EM maps, we observed that – following all-atom refinement – in incorrect regions, the models still fit the density well, but did so by introducing strain in the nearby bond angles and torsions. This often occurred in near Cβ atom of aromatic residues, where strain was introduced to fit the sidechain into density (Supplemental Figure 1). We reasoned that in these strained residues, the backbone was incorrect; by correcting the backbone we would be able to fit a non-strained sidechain into density. Thus, local strain can serve as an indicator to identify regions to refine to improve both the fit-to-density and model geometry. We developed an error predictor by constructing a function (see *Methods*) that assesses both local model-map agreement as well as local model-strain. Using a training dataset composed of error-containing models of a cryo-EM map in which the structure has been determined by X-ray crystallography (Supplemental Figure 2), we show that the new error predictor offers better discrimination of incorrectly versus correctly placed backbone, with an AUPRC (area under precision-recall curve) of 0.80 versus 0.76 using density alone (Supplemental Figure 2). In cases where models are hand-built into density, we expect this strain term to play an even larger role, as fit-to-data is expected to have larger influence on the initially constructed model.

### Better treatment of sidechain density

Recent works have shown that certain sidechains – particularly negatively charged amino acids (Glu/Asp) – tend to suffer from radiation damage and thus appear weaker in single-particle reconstructions [16,17]. Moreover, density from certain bulky sidechains, for example, Lys and Arg, tends to be less well-defined than their backbone density. This missing density dramatically affects the convergence of conformational sampling during structure refinement, where sidechains tend to be fit into density corresponding to backbone atoms. To compensate for this, we downweigh the contributions of sidechains which are less resolved in cryo-EM density. Down-weighing factors for each amino acid were determined by comparing the average per-amino-acid real-space B-factor on two cryo-EM reconstructions with known high-resolution crystal structures (20S proteasome [1] and β-galactosidase [16]), where the ratio of backbone and sidechain average B-factors was used to derive the scaling factors. Supplemental Table 1 shows the computed scalefactors used in our refinement method.

### Local sidechain refinement for large complexes

When our previous all-atom refinement approach was applied to very large complexes (800+ residues), we observed many instances where sidechains were not properly optimized into density (Supplemental Figure 3). It was hypothesized that this was due to the convergence of sidechain optimization, as the number of possible sidechain states expands exponentially with the number of residues present in a protein. Here, we opted to treat this global optimization problem as a series of smaller local optimization problems, repeatedly optimizing overlapping regions of ~20-100 residues until all residues in a protein are visited at least once. This approach resolved this sidechain fitting issue, as shown in Supplemental Figure 3 (right panel).

### Voxel size refinement

The voxel size of a cryo-EM reconstruction is determined by the physical pixel size on the detector scaled by a magnification factor. However, the magnification factor may be determined with some inaccuracy, leading to errors in deciding the voxel size of the resulting single-particle reconstruction. This is especially error prone when a reconstruction has no known macromolecular structures can be fit in to calibrate the voxel size. Here, we develop a voxel size refinement strategy, which scales the voxel size of the map to maximize model-map real-space correlation coefficient. During refinement we alternate structure refinement and map voxel-size refinement with several cycles iteratively until the voxel size converges (Figure 1). The approach is fully described in the Methods section.

### The role of independent reconstruction in structure refinement

In our previous approaches, we have used independent reconstructions (“validation” half-map) for both model selection [11] and for determination of the balance between model geometry and fit-to-data during refinement [18]. In this manuscript, we use independent reconstructions in the same manner during the first two stages of refinement (Figure 1). However, at the very last stage we perform several steps in the context of the full reconstruction, due to the additional sidechain details that may be only present in the full reconstruction. As shown in Figure 1, for the best 10 sampled models selected from the stage 2, we perform a final all-atom and atomic B-factor refinement against the complete reconstruction. Using two independent halves of the data (training/validation half-maps), the weight on the use of full-reconstruction data is optimized (see *Methods* and Supplemental Figure 3), and that weight is used in refinement against the full reconstruction. Following refinement against the full reconstruction, model geometry is verified (using MolProbity [15]) to ensure it is not worsening during refinement against the full reconstruction. This confers additional sensitivity during model selection.

### Evaluation of refined models with Molprobity and EMRinger

Models are evaluated for geometric quality using Molprobity [15], which compares summary statistics of an all-atom model to those from high-resolution crystal structures. In addition to using MolProbity to assess model quality, we further validate the Rosetta-refined models with EMRinger [19], as an independent source to validate both model geometry and density-fit at sidechain level. EMRinger samples density around C_γ_ atoms as they are rotated about the χ_1_ dihedral angle, and identifies the angle which presents peak density for the C_γ_; based on prior statistical and chemical information, this position should generally fall into the rotamer distribution of χ_1_, with angles of 60, 180, and 300 degrees. The distribution of measured peak angles at various signal-to-noise cutoffs is integrated into the EMRinger score, which reports on backbone model-to-map agreement using side chain geometry.

### Application to TRPV1

We first applied our new refinement approach to the recently determined 3.4-Å cryo-EM reconstruction of the TRPV1 channel in the apo form [12]. Half-maps were reconstructed by subdividing particles into two sets randomly, with one used for initial model rebuilding and refinement, and the other used for validation. The deposited model (PDB id: 3J5P) was used as input to the protocol described previously. All refinement was carried out using the native C4 symmetry. All input files are included as Supplemental Data File 1.

The results of refinement are indicated in Figures 2 and 3, and Table 1. The refined model improves both model quality and model-data agreement compared to the deposited model: the MolProbity score improves from 3.81 to 1.45, the fit-to-data (integrated Fourier shell correlation from 10 to 3.4Å) improves from 0.641 to 0.647, and the EMRinger score improves from 0.65 to 2.34. Figure 2A–B compares the refined model and the deposited model, colored with model violations reported by MolProbity. Figure 2C shows that our refined model – in addition to improving geometry – also improves the fit to the experimental data. Figure 3A illustrates the convergence of our refined ensemble, showing the 10 selected refined structures, the average model colored by per residue structural variation, and the refined B-factors. Both of these measures provide unique insights on assessing the local confidence of the refined models, in which structural variance shows the allowed local conformations that satisfy the density data, whereas B-factors assess the local resolution of the density data at different regions of a model.

**Figure 2.**
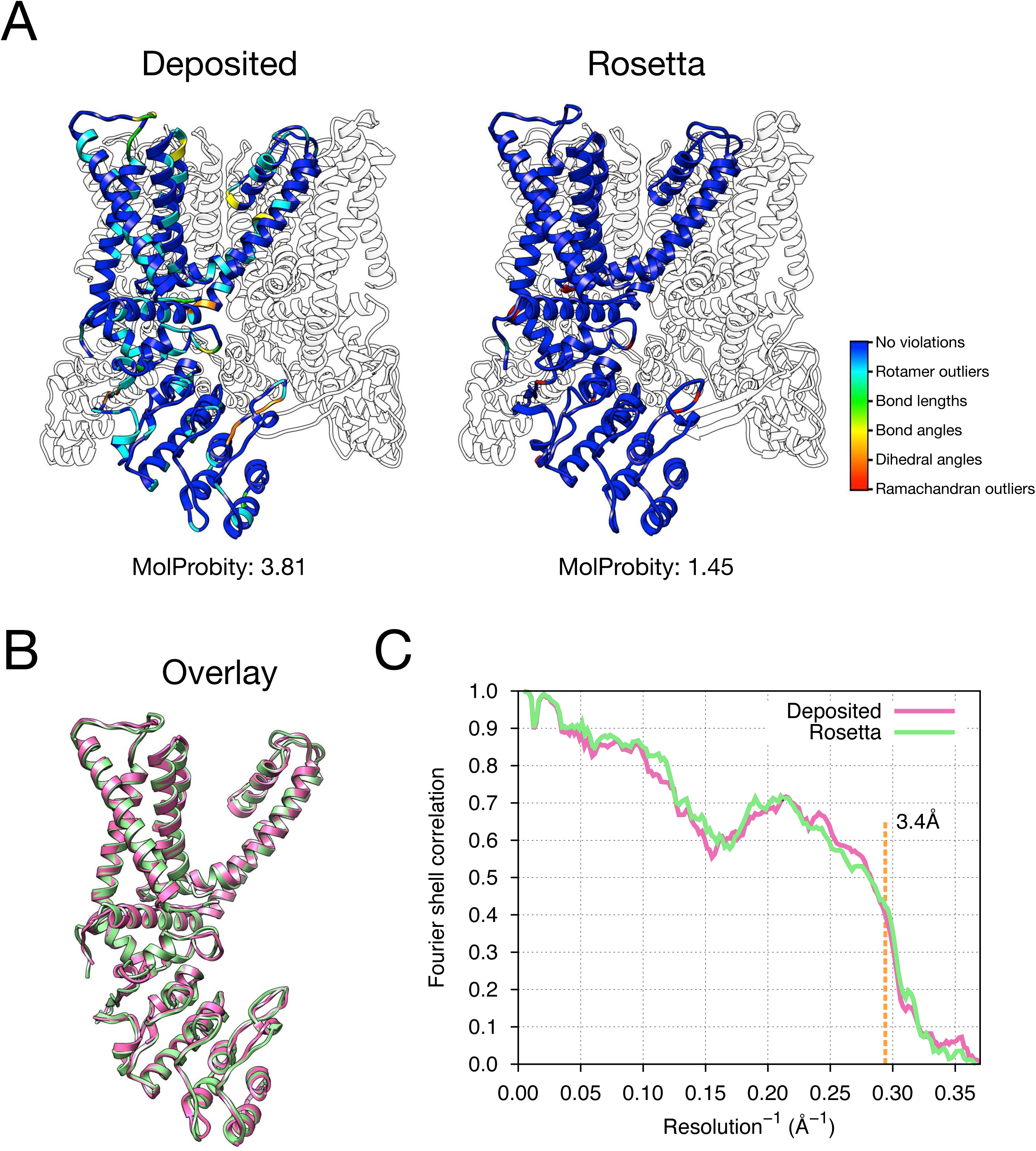
Refinement of the apo TRPV1 channel (EMD-5778) shows improved fit-to-density and model quality. (A) A comparison of the deposited and Rosetta-refined models, as assessed by MolProbity. Residues reported as violations are colored using the key shown in the far right. Blue open arrows indicate that hydrogen-bond geometry of a β-hairpin was automatically detected and improved in the Rosetta refined model. (B) An overlay the asymmetric unit of the deposited (pink) and Rosetta-refined (green) model indicates the magnitude of conformational changes that are explored by our refinement approach. (C) The agreement of models to map assessed by Fourier space correlation (Y-axis) at each resolution shell (X-axis), where the reported resolution (3.4Å) is depicted in a dashed line colored in orange. The deposited model is shown in the curve with pink color, while the Rosetta refined model is shown in the curve colored in green.

**Figure 3.**
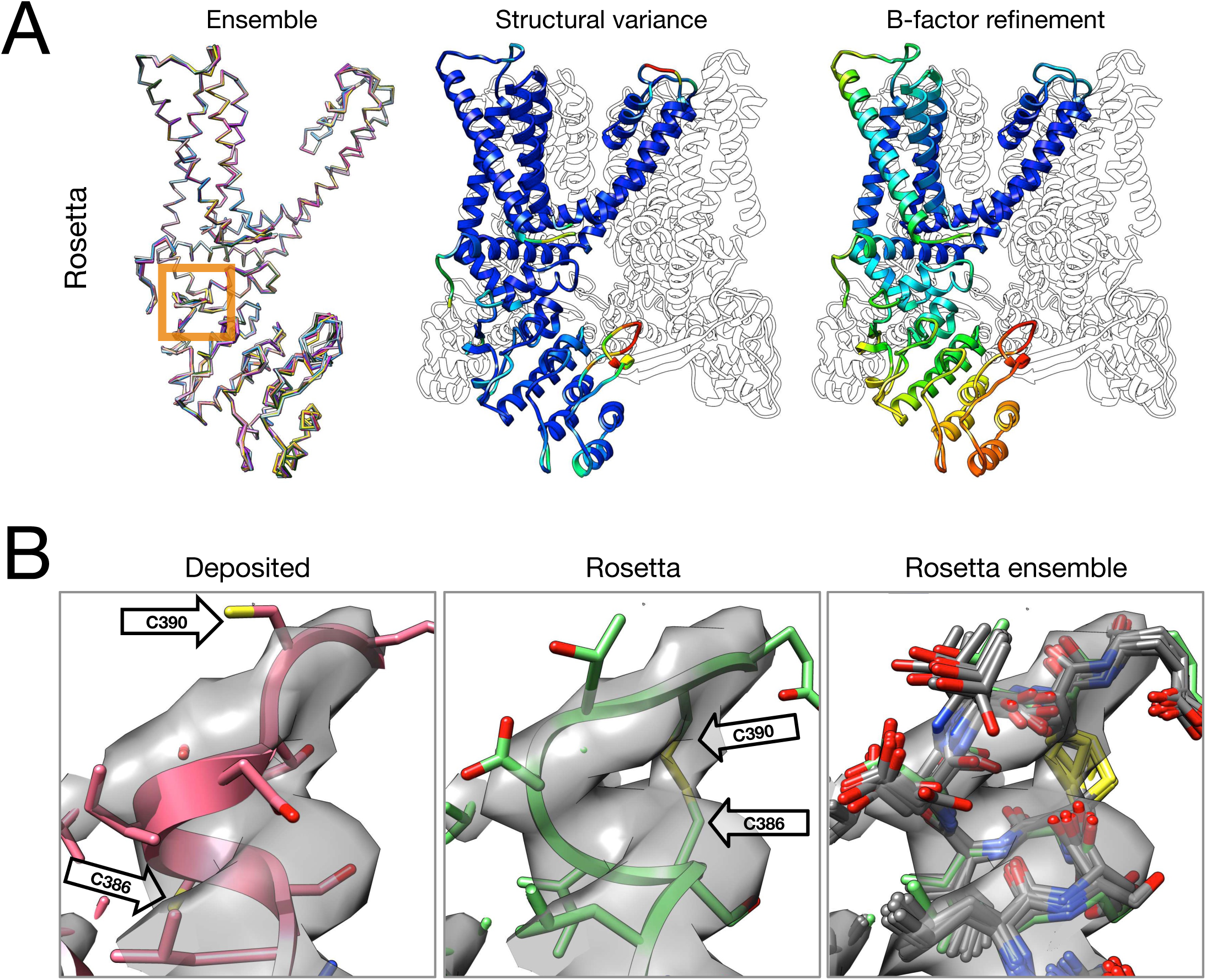
Refinement of the TRPV1 channel identifies a previously unmodelled disulfide bond. (A) An overview of the entire structure, estimating local model uncertainty in two ways: local structural diversity and refined B-factors. Local structure diversity is indicated by showing an overlay of the top 10 Rosetta models (left), the average model colored by per residue deviation (middle), and the refined per-atom B-factors (right). The orange square shows the location of a newly identified disulfide bond (C386-C390) revealed by our refinement protocol. (B) A zoomed-in view of the disulfide linkage (C386-C390) identified by the automated method. Note that the sidechain coordinates of C390 were unassigned in the deposited model; for presentation, the sidechain atoms of C390 were optimally added by Rosetta based on the deposited backbone torsion angles of C390.

**Table 1.** Structure refinement of macromolecular assemblies from cryo-EM maps using Rosetta

|  | EMD ID | PDB ID | Reported resolution [Å] | Symmetry | Number of residues<sup>a</sup> | MolProbity<sup>b</sup> |  |  |  | EMRinger Score<sup>b</sup> | iFSC<sup>c</sup> |
| --- | --- | --- | --- | --- | --- | --- | --- | --- | --- | --- | --- |
|  |  |  |  |  |  | Score | Clashscore | Rotamer outliers | Ramachandran favored |  |  |
| TRPV1 | 5778 | 3j5p | 3.4 | C4 | 489 (1956) | 3.81 / 1.45 | 86.35 / 1.96 | 28.78 / 0.00 | 95.65 / 91.93 | 0.65 / 2.34 | 0.641 / 0.647 |
| Frh | 2513 | 4ci0 | 3.4 | T | 902 (10716) | 3.97 / 1.60 | 117.06 / 3.43 | 39.11 / 0.00 | 96.51 / 92.56 | 0.31 / 2.34 | 0.659 / 0.746 |
| Mitoribosome | 2762 | 3j7y | 3.4 | N/A | 8998 | 2.71 / 1.50 | 8.38 / 3.51 | 8.49 / 0.08 | 89.86 / 94.86 | 2.09 / 2.40 | 0.734 / 0.725 |

b. Scores from deposited (left) versus (/) Rosetta refined (right) model.
c. Integrated Fourier shell correlation (iFSC) from 10–3.4Å.

a. Number of protein residues in the asymmetric unit and (the total residues) modelled.
b. Scores from deposited (left) versus (/) Rosetta refined (right) model.
c. Integrated Fourier shell correlation (iFSC) from 10–3.4Å.

Closer inspection of the refined models identified a disulfide linkage (C386-C390) that was not built in the deposited model (Figure 3B). This disulfide has previously been identified and characterized in the literature as playing an important role in response to oxidative stress for the TRPV1 channel [20]; this, combined with our models better explaining a tube of density unaccounted for in the deposited model, lets us speculate that this disulfide bond is present in the cryo-EM reconstruction. This motion also illustrates the magnitude of conformational change that may be captured by our protocol; our Monte Carlo backbone sampling strategy allows refinement to overcome energy barriers that other methods using density minimization alone cannot. Despite the magnitude of these changes, the conformational ensemble is well converged in this region (Figure 3B, right panel) providing further confidence in our refined model.

### Refinement of highly-liganded complexes: application to the F_420_-reducing [NiFe] hydrogenase complex

As our next test of the approach, we wanted to illustrate model refinement of a complex with large numbers of ligands, some of which are covalently bound, all in a system with high-order point symmetry. For this, we chose the 3.4-Å reconstruction of F_420_-reducing [NiFe] hydrogenase complex, where the asymmetric unit contains 3 protein chains which feature with a [NiFe] cluster, two metal ions, and four [4Fe4S] clusters covalently bound to cysteine sidechains, and an FAD. The complex is a dodecamer with tetrahedral symmetry, with 12 copies of a 902-residue molecule of three protein chains. We used the -*auto_setup_metals* option of Rosetta (full input files are included as Supplemental Data File 2) to maintain covalent linkages between protein and ligand during refinement. The results of refinement are indicated in Figure 5 and Table 1, where the MolProbity score improves from 3.97 to 1.60, the iFSC improves from 0.659 to 0.746, and the EMRinger score improves from 0.31 to 2.34.

**Figure 5.**
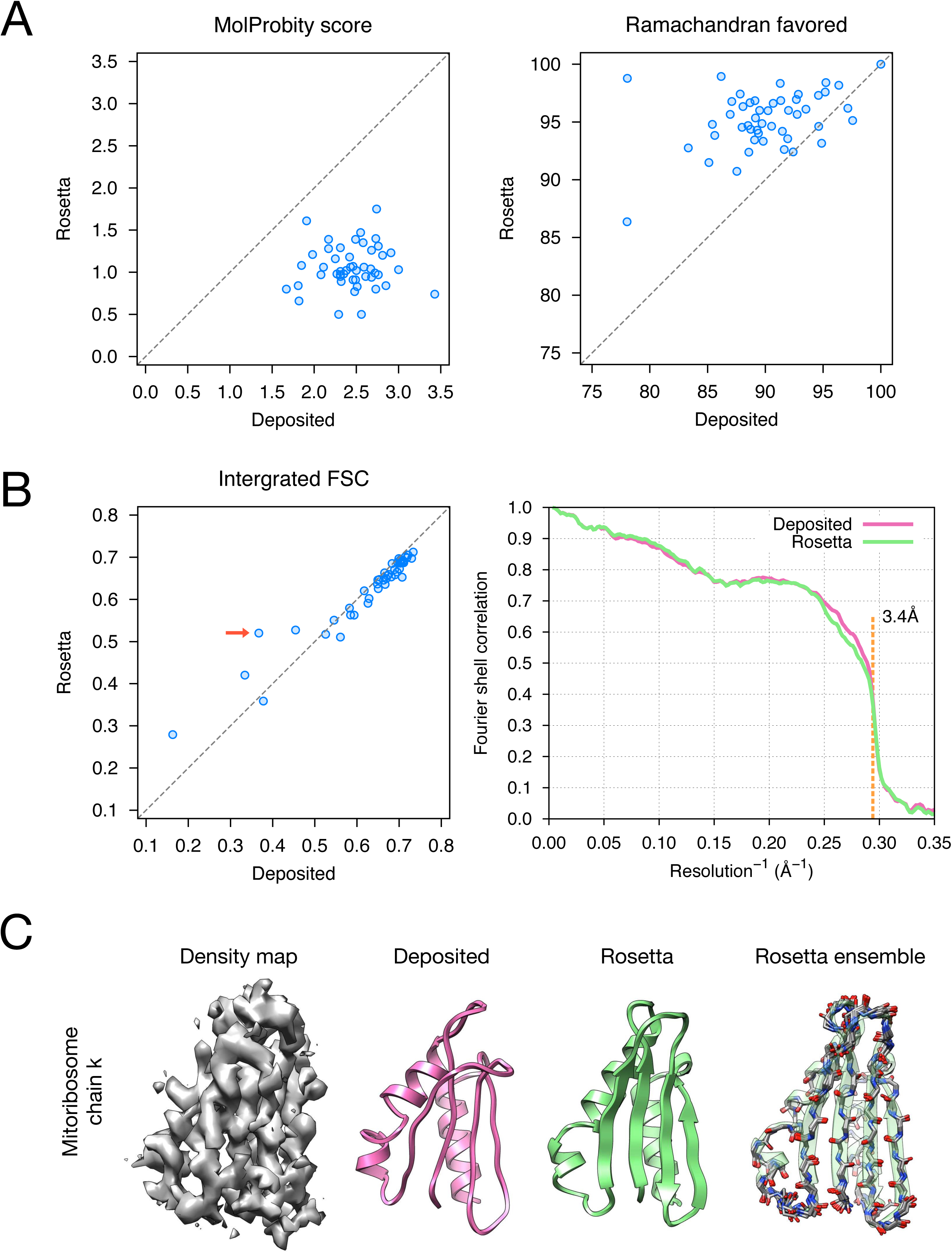
Refinement of the large subunit of the human mitochondrial ribosome (EMD-2762) shows improvements to all subunits. (A) Scatterplots of model quality of each of the 48 protein chains compare the deposited (X-axis) and Rosetta (Y-axis) models using MolProbity. On the left, the MolProbity score of all 48 protein chains are compared, where lower values indicate better model geometry. On the right, the percentage of “Ramachandran favored” residues are compared on each chain, with higher values preferable. (B) An evaluation of the fit-to-density of each protein chain. On the left, we compare the Fourier shell correlation (FSC) of each chain before and after refinement; we integrate the FSC from 10Å to 3.4Å. Higher values indicate better agreement with the data. The largest improvement, chain k, is indicated by the red arrow. On the right, the full FSC curve is shown, with the deposited model shown in pink, and the Rosetta refined model shown in green; the reported map resolution (3.4Å) is indicated in the dashed orange line. (C) A zoomed-in view indicating the large radius of convergence of the refinement of chain k. The left panel shows the density for chain k is in a region of relatively low local resolution.

Upon using EMRinger to evaluate the deposited model, we found a notably poor EMRinger score (0.31) for the symmetric complex, which contrasted with a much better score of 1.31 for its asymmetric unit (as the deposited model), indicating a high variance of density fit in different subunits. This result suggested the need for voxel-size refinement on this target. Our refinement yielded notably different voxel size than that of the deposited reconstruction: the refined model converged on 1.326Å voxel size compared to 1.320Å in the deposited map. The refined model shows better subjective agreement to the density data (Figure 4A): the deposited model clearly exposes the structure out of the density globally (Figure 4A, left panel) and locally (Figure 4A, middle and right panels), while the refined model clearly embeds its structure into the density. The refined model additionally shows better quantitative agreement to the density data (Figure 4B) and better geometry (Figure 4C): voxel-size refinement within Rosetta improved the fit-to-density and EMRinger score, compared to refinement without voxel-size refinement. The final model improved fit to data (integrated FSC from 10 to 3.4Å) by 13.2%, compared to the deposited symmetric complex (Table 1).

**Figure 4.**
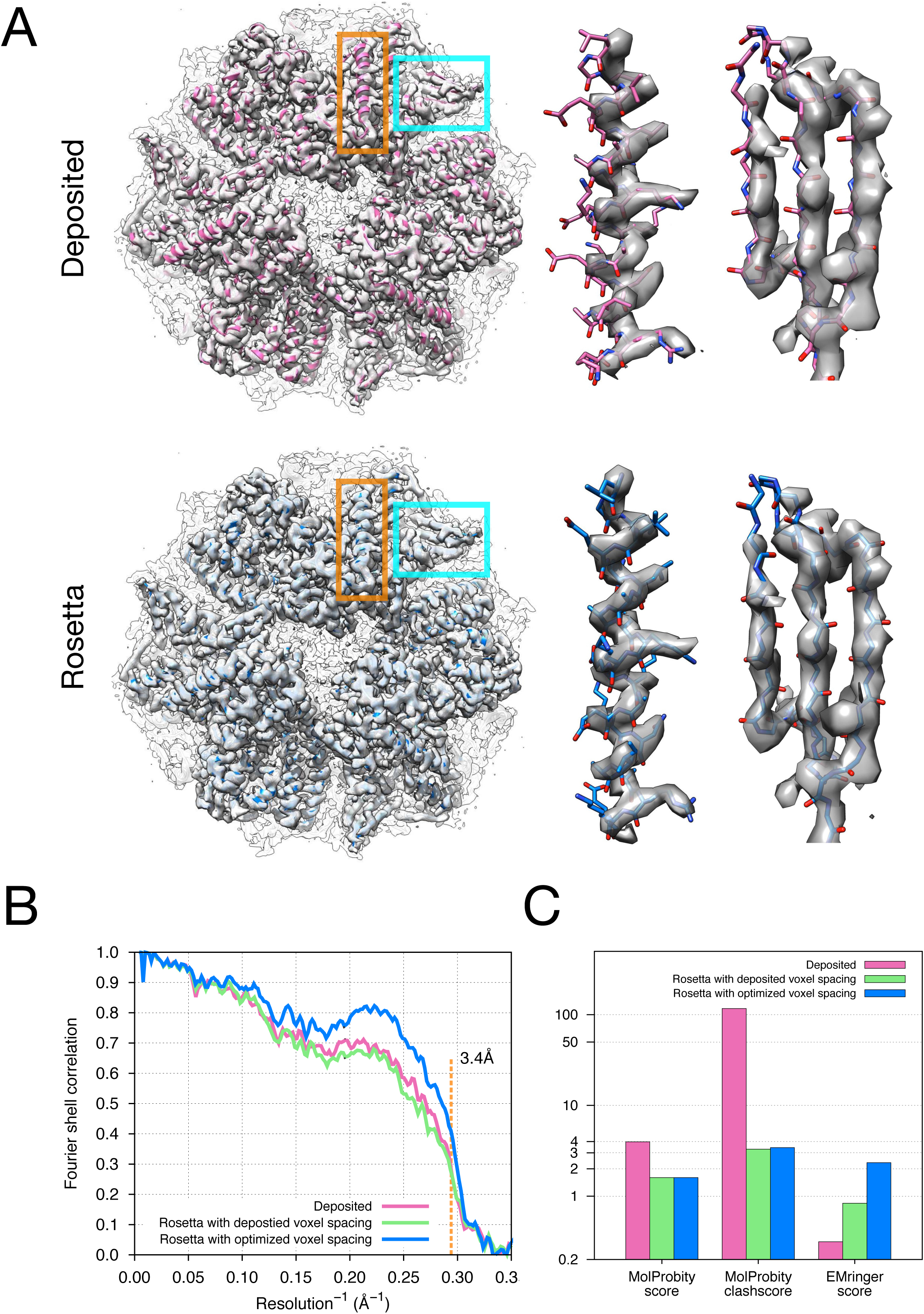
Refinement of the F_420_-reducing [NiFe] hydrogenase (EMD-2513) identifies changes to voxel size. (A) An illustration comparing the deposited and Rosetta refined models and maps. Under the same contour level (0.065), the deposited model (pink) shifts the entire complex out of the deposited density map with 1.320Å voxel size; the refined model (blue) – with 1.326Å voxel size – shows much better agreement between model and map, with much more of the model enclosed at the same contour level. The middle and left panel shows a zoomed-in view of two regions in the deposited and Rosetta refined models/maps, corresponding to the helix and the sheet indicated by the orange and cyan squares on the left panel. (B) Model-map agreement – as assessed by Fourier shell correlation (Y-axis) as a function of resolution (X-axis) – quantifies this improvement following voxel size refinement. The pink curve corresponds to the deposited model; the green curve corresponds to a model refined by Rosetta into the density map with fixed voxel size (matching the deposited value of 1.320Å); the blue line represents a model refined by Rosetta with alternate structure coordinate refinement and map voxel size optimization. (C) Model quality as assessed by MolProbity and EMRinger. The X-axis shows methods used to evaluate the models, while the Y-axis shows the scores under each criterion.

### Refinement of large complexes: application to the mitochondrial ribosome large subunit

Finally, we wanted to test the ability of our refinement to scale to large asymmetric macromolecular assemblies, more typical of cryo-EM single particle reconstruction. To do so, we considered refining models against the previously published 3.4-Å cryo-EM reconstruction of the large subunit of the human mitochondrial ribosome [14]. The deposited model had been previously refined with *REFMAC* [9], and consists of 48 chains with 8998 amino acids assigned and 1628 nucleic acid bases.

In order to make conformational sampling tractable, we used a slightly modified strategy from that shown in Figure 1 (full input files are included as Supplemental Data File 3). The first two steps of the protocol (error identification and backbone rebuilding) were carried out on each protein chain individually, while the third step was carried out on the fully assembled complex. Model selection was carried out on each individual chain; each selected model was refined as a complete assembly, with the top model of each chain refined together, the second selected model for each chain refined together, and so on. Nucleic acids were not refined but were included as rigid bodies to accurately recapitulate protein/RNA interactions.

The results of refinement are indicated in Figure 5 and Table 1. Several large-scale conformational changes again appear in converged models; these models show better geometry, fit to density and fewer unexplained regions of density. The backbone geometry improvements are in particular noticeable in proteins with β-sheet containing domains. Unlike other refinement procedures (*phenix.real_space* [10] and *REFMAC* [9]), which require manual input of secondary structure restraints determined either from an initial model or homologous protein structure to maintain backbone geometry during refinement, in our approach the Rosetta forcefield is able to optimize hydrogen bond geometry in secondary structures without requiring *a priori* knowledge of secondary structures. This is particularly powerful in refining *de novo* structures where secondary structure is ambiguous due to poor local resolution. Figure 5C illustrates an example (chain k) of this from the case of mitoribosome, where a β-sheet not present in the original model is identified, the backbone geometry is improved, and the model fits the density much better than the deposited model (Figure B, left panel, red arrow); the refinement also shows a large radius of convergence.

The refined model ribosome model has 1.50 MolProbity score, 0.725 iFSC, and 2.40 EMRinger score. The largest improvements tend to occur in regions of low local resolution (~5Å assessed by ResMap from the original paper) on the periphery of the complex. Looking at the results on individual chains, as indicated in Figure 5A and Supplemental Table 2, the MolProbity score improves on all 48 protein chains, which in part is from the much improved backbone geometry assessed by the *Ramachandran favored* term in MolProbity (Figure A, right panel and Supplemental Table 2). Our Monte Carlo backbone sampling can correct these incorrect backbone placements, which often require significant compensating conformational changes. EMRinger score is also consistently improved (Supplemental Figure. 4), particularly in regions where the deposited model scores poorly.

## Discussion

In this manuscript, we develop a method for improving atomic details of manually traced models from 3-4.5Å resolution cryo-EM density. We show the applicability of the approach, by applying it to three systems: a membrane protein, an asymmetric macromolecular assembly containing large numbers of protein chains and RNAs, and a highly symmetric system with a large number of ligands. In all cases, we show that we are able to significantly improve model geometry while maintaining or improving agreement to the density data. We show that model convergence can be used to suggest local model uncertainty in addition to B-factors. Finally, we also show that our models also recover structure features that are supported in the literature, or in much better local agreement with the density data.

Unlike other approaches [9,21], our approach can automatically perform large-scale backbone reorganization, correcting backbone placement errors common in these 3-4.5Å resolution datasets. Two features of our refinement approach regarding the use of prior information are critical in the success of this large-scale refinement. First, the use a physically realistic forcefield throughout refinement handles the under-constrained nature of refinement at these resolutions, by using chemical “domain knowledge” learned from high-resolution crystal structures to implicitly fill in the missing information in the data. Second, our fragment-based rebuilding which explicitly samples the most likely backbone conformations given a short stretch of sequence also uses prior information gather from high-resolution protein structures, further restricting conformation space, and filling in additional information not present in the data.

Finally, an open question is on what way structure refinement can be further improved, particularly as refinement extends to even lower resolutions (worse than 5Å). Enhancing the predicting power of the Rosetta modeling methods is the key to to push the resolution limit of the current refinement method further. This can be achieved through: 1) improving the energy function (forcefield) used in refinement, and 2) improvements in conformational sampling methodology, particularly for systems where secondary structure prediction is poor. Further improvements in the role of B-factor sharpening and the effect on refinement are necessary, as well as better predictors of local model error. Finally, structure refinement in maps with highly heterogeneous local resolution remains challenging, where a single set of refinement parameters cannot readily be applied at all regions. Methodological improvements that that allow adjustment of parameters based on local map quality will be essential to accurately refine structures from such maps. In our effort to enable automated structure refinement on large macromolecular assemblies, we hope this method can be a valuable tool for determining atomic accuracy structures from near-atomic-resolution cryo-EM data.

## Methods

### Preparing maps for refinement

Split maps were provided by the original authors. One map was randomly chosen for refinement, and the other was used for validation. In all, cases a B-factor of −100 was applied to the map used for refinement using the “*image_handler”* tool in RELION [3]. The maps were subsequently filtered to the user-refined resolution. In the case of the mitochondrial ribosome, segmented maps were prepared using a custom Rosetta application and the deposited structure to guide segmentation:
*density_tools.default.linuxgccrelease-s 3j7y0.pdb-mapfile EMD-2762.mrc-mask_radius 2-maskonly*

Some steps of the protocol also made use of the full reconstruction. As with the training map, these were sharpened using a B-factor of −90 with a low-pass filter to 3Å.

### Preparing structures for refinement

In the case of TRPV1, residues 111-202 in the Ankryin repeat domain from the deposited model did not have visible density, and so were delete proior to refinement. Furthermore, automatic refinement as applied in two stages due to the highly heterogeneity between the trans-membrane domain and the Ankryin repeat domain. The trans-membrane domain (residue 234-586) was first refined in the density masked using the deposited model. In the case of the mitoribosome, residues from chain t and chain f, in which atoms are assigned to residues “UNK,” were removed from all the refinement process, as well as data analyses or results comparisons. In the case of Frh, refinement of ligands received special treatment: refinement started using protein only, with constraints maintaining ligand site geometry. Later, ligands were added back on and rerefined.

### Algorithm for model rebuilding

Model rebuilding generally follows the procedure from our previous work [11], with a few key changes highlighted below. Rebuilding starts from the deposited structure, which is first conservatively refined using one macrocycle of the Rosetta *relax* protocol to trigger local strain on sidechains, which iterates four cycles Monte Carlo rotamer optimization with all-atom minimization, ramping the weight on van der Waals repulsion in each cycle. Minimization is carried out in Cartesian space, with a term enforcing ideal bond angles, bond lengths, and planarity [22].

Following Cartesian minimization, the worst residues are selected using the following equation to evaluate the quality of the model at residue *i*:

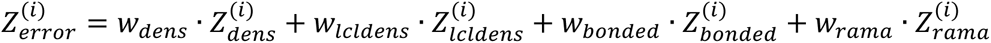

Four different terms appear in this equation, two of which assess a model’s agreement to data, two of which assess a model’s local strain. The first two, 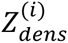 and 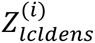, assess the model-map agreement of the backbone and sidechain atoms of each residue, computing the real-space correlation coefficient in a region around a residue, and converting that to a *Z-score* compared to the entire model. For the former term, an absolute correlation coefficient is computed; for the latter term, the correlation is normalized with respect to residues nearby (those within 10 Å of residue *i*). The latter term is specifically added to deal with maps that have significant diversity in local resolution.

The second two terms, 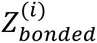 and 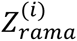, assess a model’s strain following model refinement. The motivation for these terms is that in cases where the model was built incorrectly into density, it will be energetically unfavorable. Following an initial refinement, these incorrect portions will either be move away from the data, or will introduce model strain to maintain the favorable agreement to the data, depending upon the balance of forces between the two. These terms compare the per-residue bond geometry term, and the per-residue Ramachandran energy, respectively, to that over the entire structure, and compute a *Z-score* for each residue.

For each of the four terms, a Z-score is computed and is summed together, with a particular weight for each term. The weights were tuned using a 3.3-Å cryo-EM map dataset with known high-resolution structure (the 20S proteasome [1]), where a set of ~500 error-containing models was used as the training data. The results of this tuning process are shown in Supplemental Figure 1. The final weights selected were *w*_*dens*_=0.45, *w*_*lcldens*_=0.05, *w*_*bonded*_=0.15, *w*_*rama*_=0.35.

After computing this weighted Z-score for each residue, all residues with a score below some target value (see the next section on iteration for specific values) are selected for local rebuilding. Local rebuilding uses the iterative fragment-based approach previously published [11]. In our new approach, a residue is randomly chosen from the pool tagged for rebuilding from the previous step. Given the local sequence around this selected residue, a set of 25 protein backbone conformations from high-resolution structures with similar local sequence and predicted secondary structure is sampled. Each sampled backbone is refined – as an isolated fragment – into density using the following three step procedure: a) the backbone only is minimized in torsion space using a simplified energy function, b) sidechain rotamers are optimized into density, and c) both backbone and sidechain are minimized in torsion space using a simplified energy function. Constraints on the ends of each fragment ensure the local region is reasonable in the context of the entire backbone. Of the 25 sampled fragments, the best is selected by fit to density. Finally, the replaced fragment is minimized in the context of the complete structure. This process is run as a Monte Carlo trajectory.

### Iterative rebuilding and all-atom refinement

Model rebuilding and all atom refinement are run iteratively, as shown in Figure 1. Four separate 200-step Monte Carlo trajectories are run with increasing coverage of predicting errors but sacrificing the accuracy of the predictions. This is done with the Z-score cutoff increased in each step, following the schedule shown in Supplemental Figure 2: first residues with *Z*<-0.5 are selected for fragment-based rebuilding, followed by −0.3, −0.1, and finally *Z*<0. Between each cycle, a single iteration of *Relax* is run, in the same manner as the pre-refinement step. At the start of each stage, 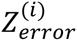 of a model is re-evaluated as above to avoid refining fixed errors from the previous stage, and residues predicted to be in error are selected. Finally, an additional 200 step Monte Carlo trajectory is run with the 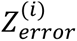 computing solely from 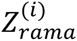 to ensure the favorable Ramachandran geometry in models.

### Pre-proline Ramachandran potential

Following early experiments, a new term was added to Rosetta that enforces a distinct pre-proline Ramachandran potential, replacing the original 20 different potentials:

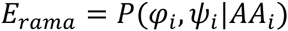

With 40 different potentials conditioned on the sequence identity of the C-terminal adjacent residue:

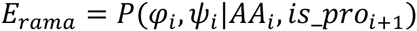

This potential was trained using the Richardson 8000 set of high-resolution crystal structures [15], and smoothed using adaptive kernel density estimates, as with the original Ramachandran potential [23]. They are included in the released Rosetta with the energy term *rama_prepro* (using the same weight as the Rosetta term *rama*). Supplemental Figure 6 illustrates the resulting potentials. For all experiments in this manuscript, this term replaced the default Ramachandran score term in Rosetta.

### Local relax

Following our four cycles of refinement, we run a modified version of *Relax*, which we call *LocalRelax*. Modifications were made following the observation that – when applied to very large complexes (800+ residues) – we observed many instances where sidechains were not properly optimized into density, even though the density was very clear. Supplemental Figure 3 shows several such cases.

In *LocalRelax*, small overlapping regions of ~20-100 residues (discontinuous in sequence space) are selected for optimization repeatedly, until the entire protein has been optimized at least once. The approach is based upon the idea of neighbor residues, where residue neighbors are defined as all residues with a Cβ-Cβ distance less than 8Å. We first find the residue *r*_*i*_ with the most residue neighbors. Then we optimize the neighbors of *r*_*i*_, and the neighbors-of-neighbors of *r*_*i*_: the neighbors are allowed to optimize both sidechain and backbone conformation, while the neighbors-of-neighbors may only optimize sidechain conformation. This optimization is performed via Monte Carlo sampling of sidechain rotamers, followed by Cartesian minimization of all movable atoms. Following this, all neighbors of *r*_*i*_ (as well as *r*_*i*_) are marked as visited, and the process repeats, selected a new *r*_*i*_ as the unmarked residue with the most neighbors. This process continues until all residues are marked. In total, 4 cycles of this procedure are carried out, increasing the weight on van der Waals repulsion in each cycle. Finally, following coordinate refinement with *LocalRelax*, we fit atomic B-factors following the scheme of our previous paper [11].

### Sidechain rescaling

We compute a scalefactor associated with each sidechain, that describes how much contribution to the density score each sidechain contributes. The values were computing using the 3.3-Å reconstruction of the 20S proteasome [1] and the 3.2-Å reconstruction of β-galactosidase [16]. Models were refined into the density and real-space atomic B-factors were fit for each atom. We then converted the atomic B-factors to scale factors using the following transformation:

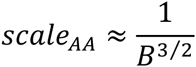

Scales were normalized such that the scale for all backbone atoms was equal to 1. To prevent overfitting, each sidechain was grouped into one of three classes, and all sidechains within a given group were given the average scalefactor of the group. Finally, while maintaining the ratio of these three groups with respect to one another, we scaled the relative contribution of backbone versus sidechain density, and selected the best values based on free FSC following refinement. The final values range from 0.66 to 0.78, and are tabulated in Supplemental Table 1.

### Voxel size refinement

To optimize the voxel size of a map used to refine the model, we fix the model coordinates, and compute the model density. We then refine the voxel size *v*=[*v*_*x*_,*v*_*y*_,*v*_*z*_] and the origin *o*=[*o*_*x*_,*o*_*y*_,*o*_*z*_] of the map density – fixing these parameters in the model density – to maximize the real-space correlation coefficient between the two:

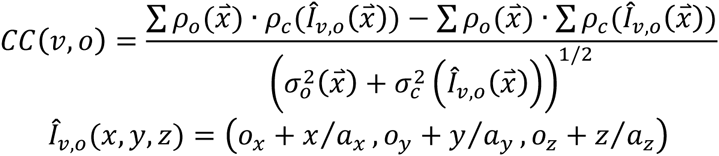

Here, *ρ*_*o*_ refers to the experimental map and *ρ*_*c*_ to the map derived from the model, while *σ*_*o*_ and *σ*_*c*_ refer to the standard deviations over the corresponding density maps. Sums are taken over the entire map. Off-grid density values are computed using cubic splines to interpolate the calculated density map. This function is optimized with respect to the voxel size paramters using l-BFGS minimization; analytic derivatives are computed for CC with respect to *v* and *o*, and the same cubic splines are used to calculate derivatives with respect to the calculated map.

Voxel size may be refined isotropically or anisotropically (either 4 or 6 total parameters); all experiments in this manuscript treated this refinement isotropically (that is, all three axes are scaled together).

### Refinement against the full reconstruction and model selection

The previously described protocol was run to generate 5000 independent trajectories. From these 5000 models, a set of 10 representative models is chosen, following the protocol outlined in Figure 1. We want our optimized models to simultaneously be optimal in terms of: a) independent map agreement, b) physically realistic geometry, and c) agreement to the full reconstruction. The latter is necessary, as the full reconstruction often features details not present in the independent half maps.

Independent-map FSCs were computed against the validation map – subject to the same sharpening scheme as the training map – using the *ComputeFSC* mover in Rosetta. The integrated FSC between 10Å and the reported resolution (3.4Å in all cases) of the map was used to assess agreement with the independent map. The script computes FSC after masking the map with a mask computed from the model and filtered to 12Å with the command line:
*density_tools.exe-in:file:s model.pdb-mapfile validation_map.mrc-mask_radius 12-nresbins 50-lowres 10-hires 3.4-verbose*

In the case of the mitochondrial ribsosome, each segmented domain map was evaluated separately. Of the 1000 generated models, the top 50 by independent map agreement are selected.

Next, we want to identify the models from this subset that are the most physically realistic. To do this, all 50 models are rescored with MolProbity [15], and the top 10 are selected. While computing similar features to the Rosetta energy, its slightly different implementation makes it a somewhat orthogonal measure for structure evaluation.

Finally, we want to use features from the full reconstruction to further improve the model, particularly bulky sidechains that may not be visible in the half-map reconstructions. However, when refining against the full reconstruction we need to be careful not to overfit to the full reconstruction, as we no longer have an independent map with which to evaluate overfitting. We use two ideas to avoid overfitting in this case. First, we do not perform any fragment based rebuilding with the full map, and instead only perform two cycles of *LocalRelax* and B-factor refinement with the full map. Second, we use halfmaps to determine the optimal fit-to-density weight when refining against the full map. The weight is selected using the following relation where the weight is chosen to maximize the following:

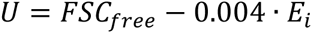

Here, E_i_ is the per-residue energy, and is included as additional regularization to avoid overfitting. The value of 0.004 was chosen to normalize the two based on the relative dynamic ranges of both terms.

The top 10 models from the previous selection are subject to refinement against the full map. The final model is then taken as the model with best integrated-FSC against the full reconstruction. Local deviation over all ten models is used to estimate model uncertainty. The per-residue structural variance of ensemble models is calculated using Theseus with the default command line [24].

### Assembly of the mitochondrial ribosome

In the case of the mitochondrial ribosome, we refine separate models for each protein subunit. A final assembly step combines the full model. In this final assembly step, all subunits, plus the deposited nucleic acid chains are combined in a single model, and are subject to 2 cycles of *LocalRelax* against the full reconstruction.

### EMRinger score calculation

For each of the five models following model selection, EMringer was run using the command:
phenix.emringer MODEL.pdb MAP.ccp4

To calculate per-chain EMRinger scores, pdb files were first segmented by chain ID and then emringer scores were calculated against the segmented pdb files. A script is included to automate the PDB segmentation and calculation of EMRinger scores.

EMRinger scores can be compared absolutely between structures, although model size and local resolution variation are sources of noise for the EMRinger score. Scores below one are indicators of suboptimal model to map agreement for structures better than 4-Å resolution, while a score around zero indicates no improvement beyond randomness.

## Availability

All methods described are available as part of Rosetta, using weekly releases after week X, 2016. The Rosetta XML files and flags for running all the refinements discussed in this manuscript are included as Supplemental Data Files 1-3. The scripts and the tutorial used for running the method described here is available now at the website of the corresponding author (https://faculty.washington.edu/dimaio/files/density_tutorial_sept15_2.pdf).

## Acknowledgements

The authors thank Drs. Alan Brown, Alexy Amunts and Venki Ramakrishnan for sharing the half maps of mitoribosomal large subunit (EMD-2762) with us; the author especially thank Dr. Alan Brown on providing helpful comments on the Rosetta refined mitoribosome, in which the suggestions led to the new development on better optimizing sidechains in very large protein complexes. The author thanks Dr. Metteo Allegretti and Janet Vonck for sharing the half maps of Frh (EMD-2513) with us. The authors thank Dr. Erhu Cao for commenting on the refined TRPV1 model initially. The authors thank Dr. Vikram Mulligan for helping on using the “-*auto_setup_metals*” module he developed on facilitating ligand setup in modeling Frh using Rosetta.

## Contributions

R.Y.-R.W. performed the research. R.Y.-R.W. and F.D. developed the method and prepared the manuscript. R.Y.-R.W., Y.S. and F.D. conceived and designed the research. F.D. supervised the research. B.A.B and J.S.F. performed the EMringer analysis. Y.C. provided the TRPV1 half-map data set with various b-factor sharpening, and analyzed and interpreted the results on the TRPV1 refinement. All authors edited the manuscript.

---

**Supplemental Figure 1.**
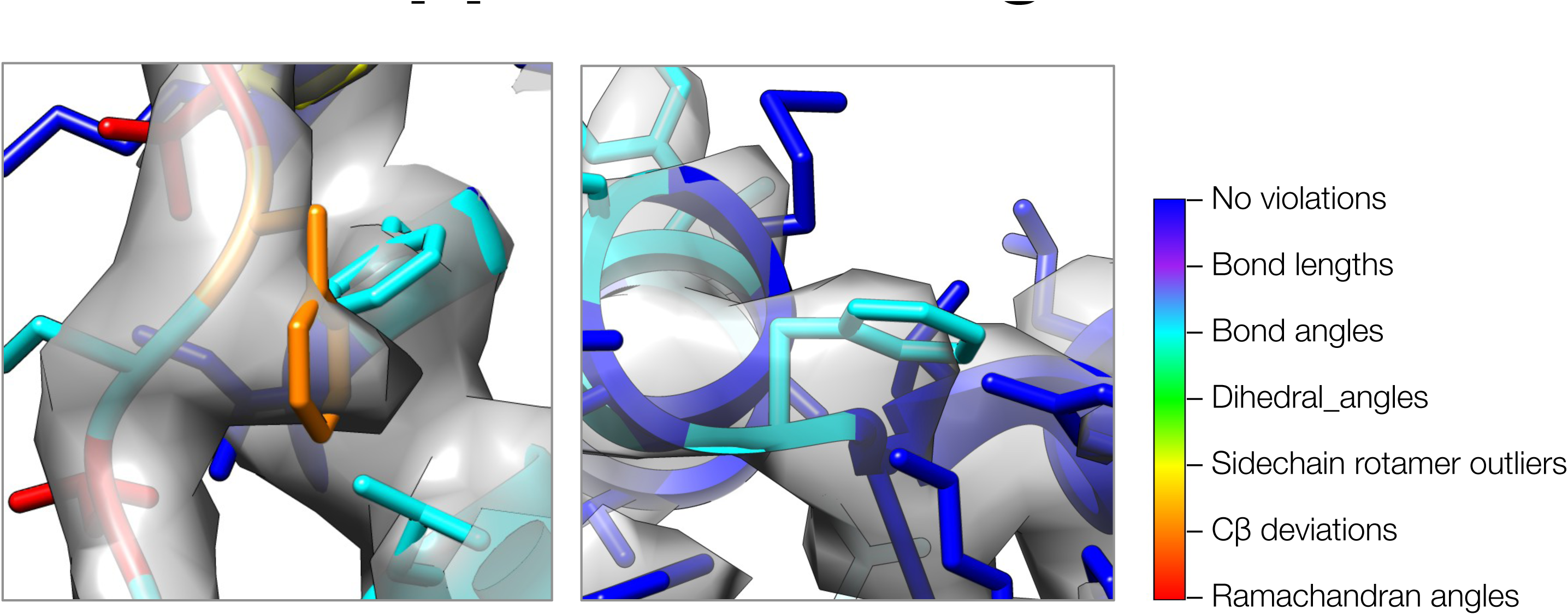
A closeup view of model strain indicating errors in density-optimized TRPV1 models using the previous Rosetta approach. Both insets show two regions of models refined by the previous approach, where strain can indicate errors in models. In both cases, phenylalanine sidechains fit the density well, but both show geometric strain around the C**β** atom. The type of strain (as evaluated by MolProbity) is indicated by model color, using the key on the right.

**Supplemental Figure 2.**
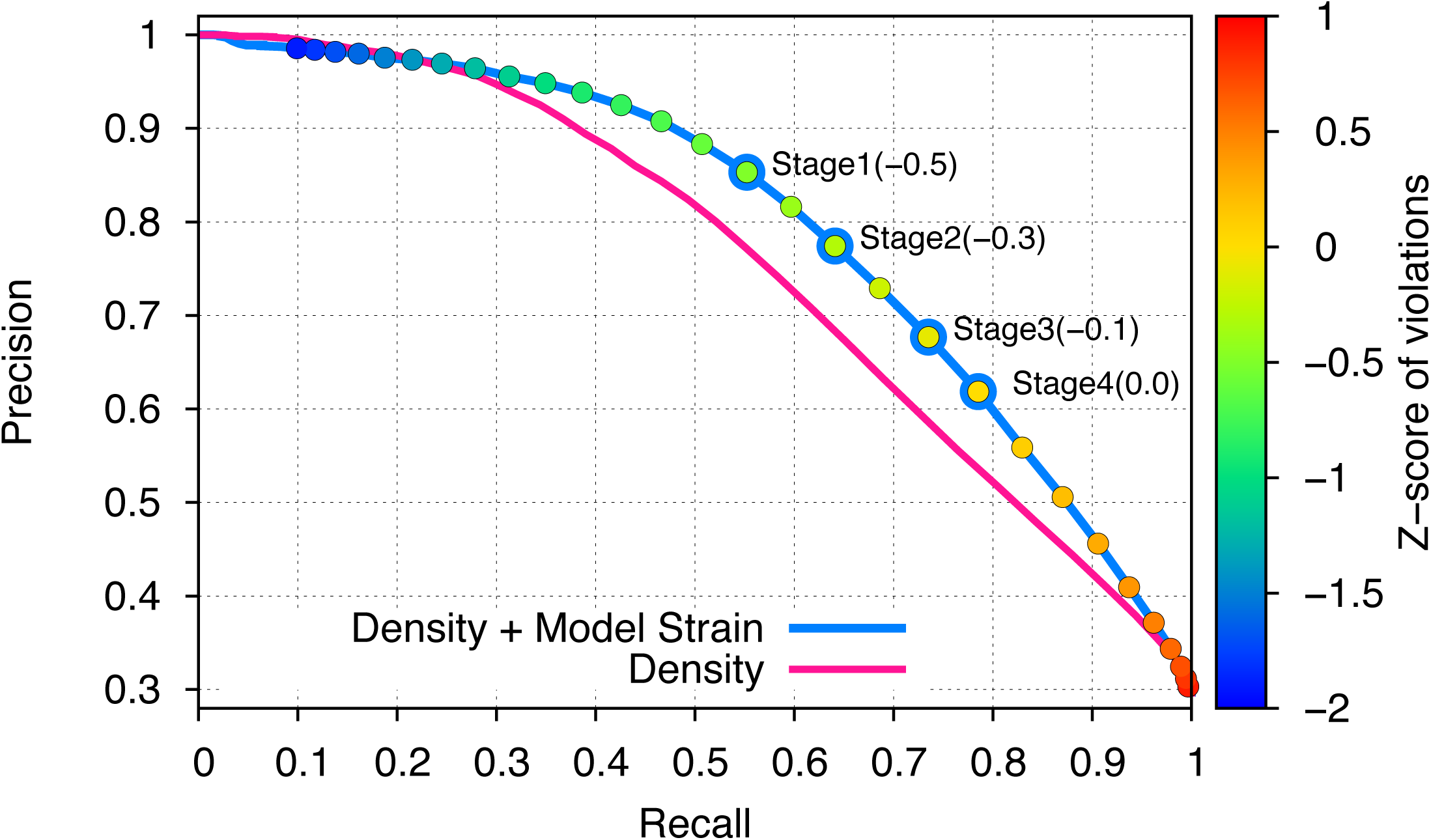
Incorporating model strain improves error detection. Guided by the 3.3-Å 20S proteasome reconstruction, we evaluated 500 models against the high-resolution crystal structure. We plot here the precision (y-axis) and recall of predicting which residues were incorrectly placed (RMS > 1Å). Using density alone (pink line) is outperformed by using a combination of density and model strain (blue line). Our refinement approach considers four points on this curve when picking density + model strain cutoffs, indicated on the plot with “Stage1-4”.

**Supplemental Figure 3.**
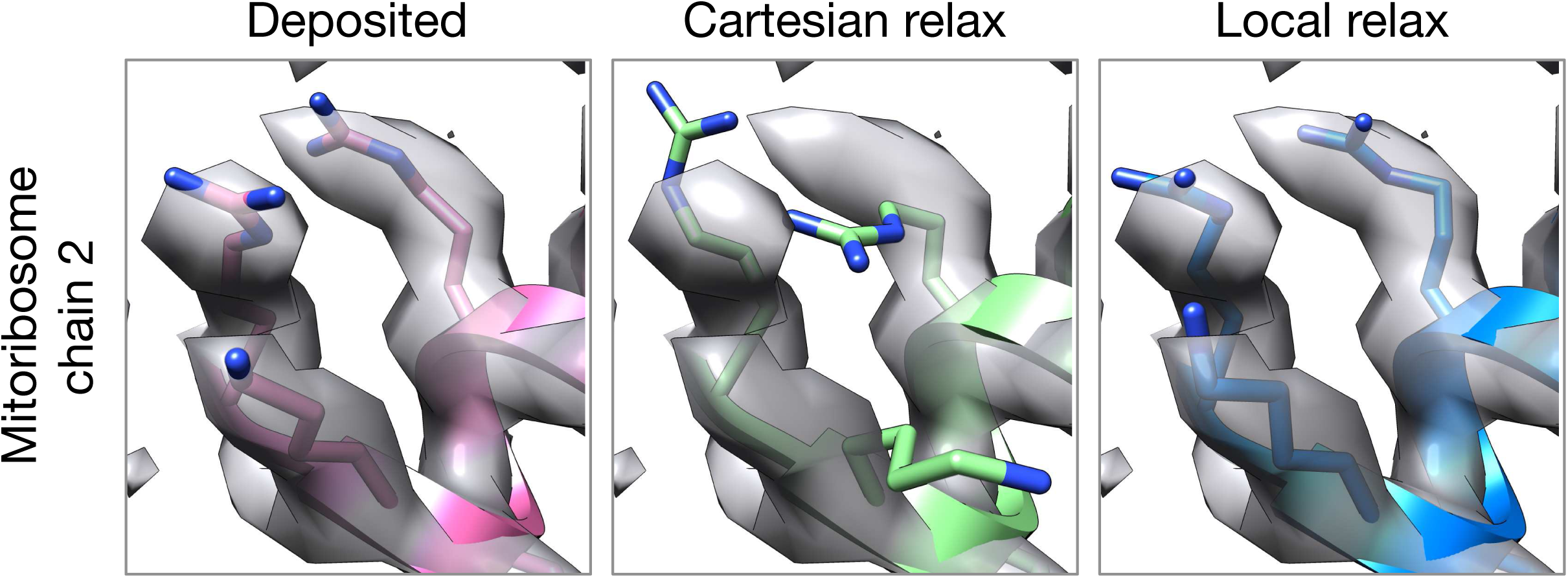
Local relax shows better placement of sidechains for large systems. In the case of mitoribosome, refinement of a particularly well-resolved region in the map (left) led to sidechains clearly misaligned with the density (middle). This was due to the poor convergence of our Monte Carlo sidechain placing approach when applied to systems with >1000 residues. Our alternative approach, LocalRelax, which instead performs many local sidechain optimizations, correctly places sidechains consistent with density (right). **Supplemental Figure 3. Density weight optimization against halfmaps for Mitoribosome**. Before refinement against the full reconstruction, we optimize the weight on the “fit-to-density” energy using half maps, to avoid overfitting. We plot several key metrics here as a function of weight on the fit-to-density score term (X-axis), including the FSC “overfitting” (FSCwork - FSCfree, top), the Rosetta energy (row 2), and several Molprobity model geometry terms (rows 3-6). In all cases, we see a sharp inflection point where overfitting increases and geometry gets notably worse. As a general rule-of-thumb, we use the weight maximizing FSCfree-0.04*per-residue-energy to capture this inflection point.

**Supplemental Figure 4.**
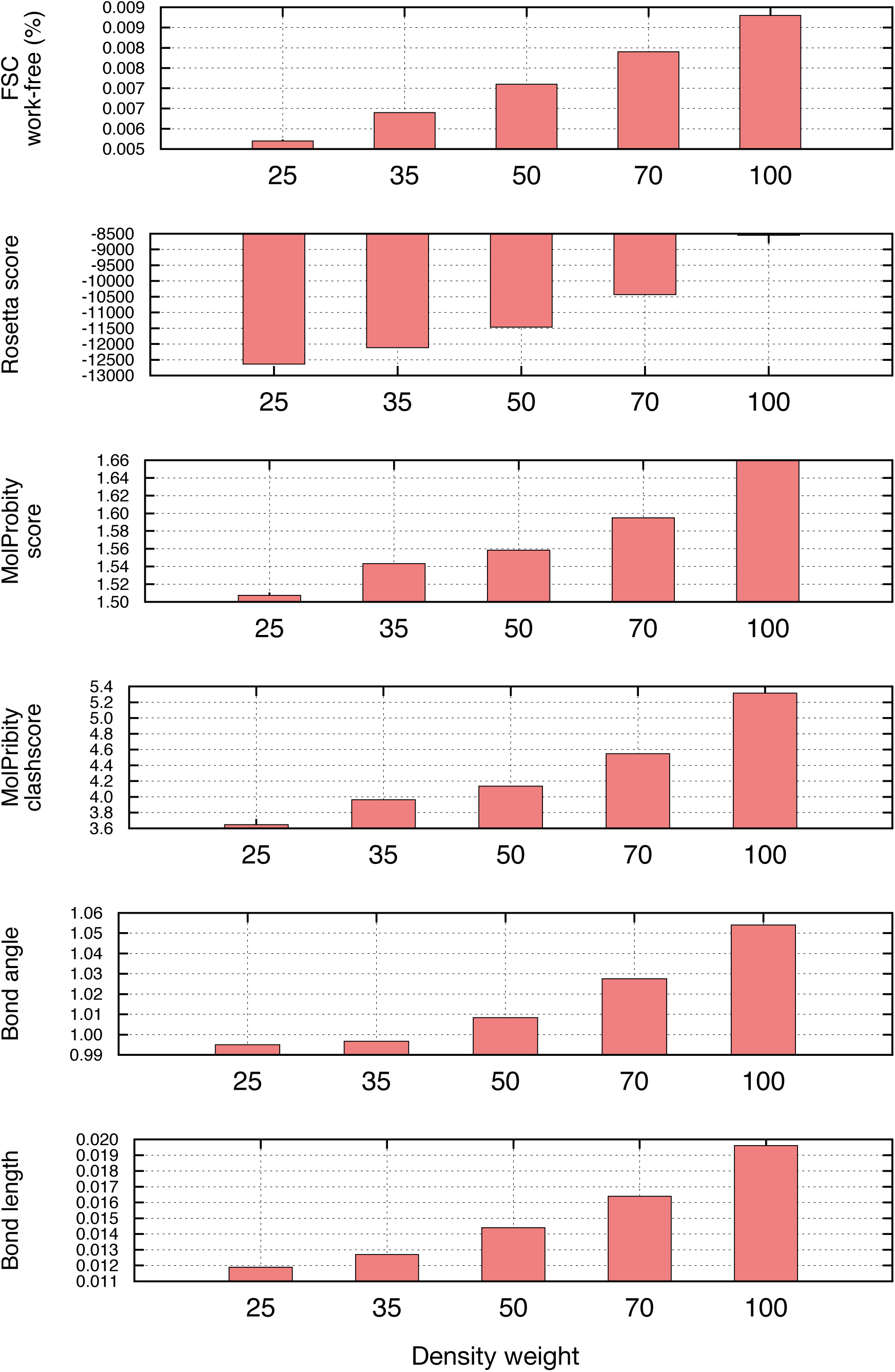
EMRinger analysis on refinement of the large subunit of the human mitochondrial ribosome. A scatterplot of model quality assessed by EMringer of each of the 48 protein chains compares the deposited (X-axis) and Rosetta (Y-axis) models.

**Supplemental Figure 5.**
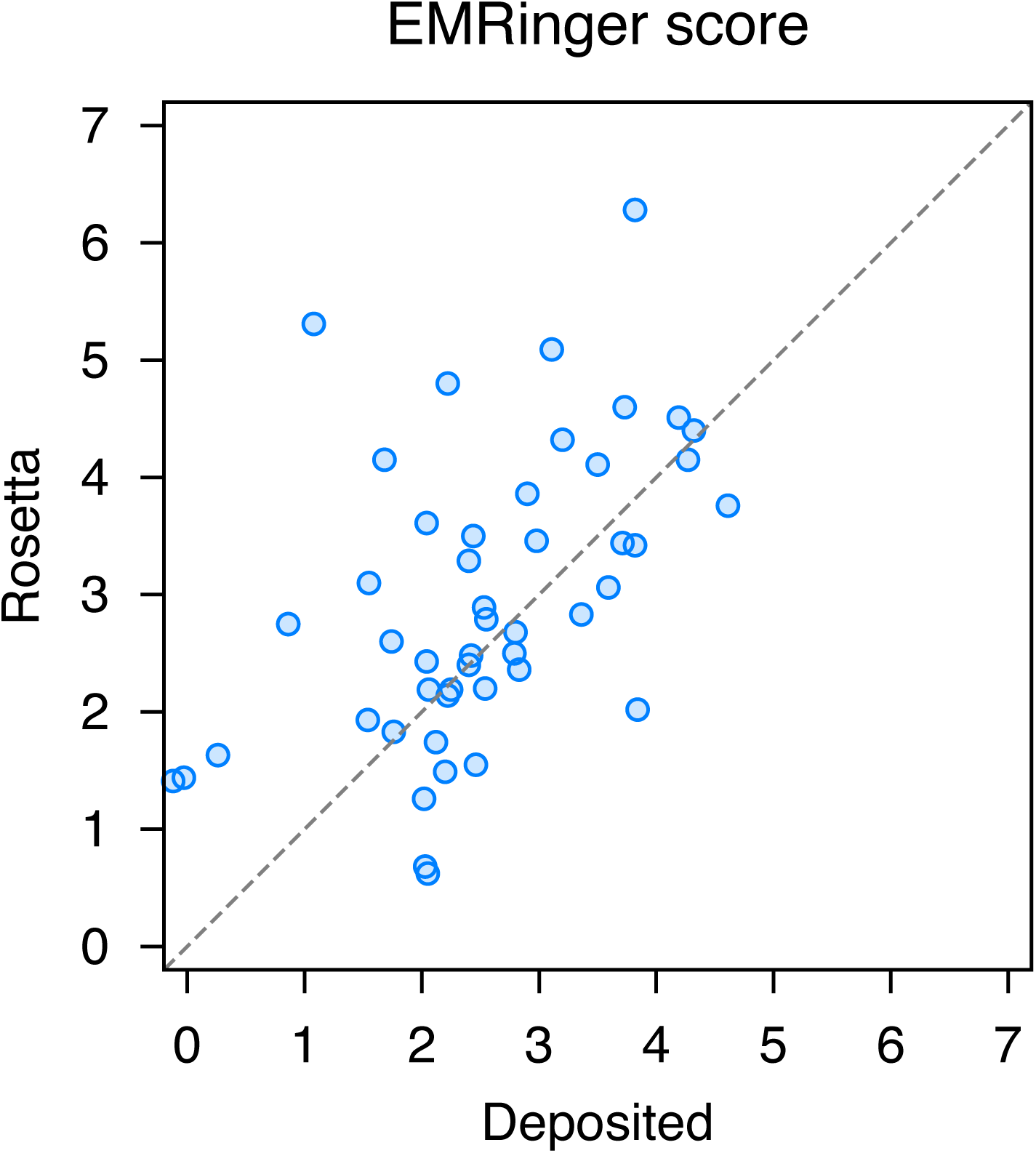
Model geometry is improved with a separate pre-proline potential. It was found that refined models initially had poor pre-proline geometry. Thus a new backbone torsional potential was created which separately treats pre-proline and pre-non-proline residues. In the plot above we show the old potential (left), the new pre-non-proline potential (middle), and the pre-proline potential (right), for three different residue identities. The color indicates the unweighted energy values, using the key on the right.

**Supplemental Table 1.** Sidechain scaling factors used in automated Rosetta structure refinement

| Sidechain | Raw data | Factor<br>used |
| --- | --- | --- |
| ARG | 0.84 | 0.66 |
| LYS | 0.84 | 0.66 |
| GLU | 0.85 | 0.66 |
| MET | 0.87 | 0.66 |
| ASP | 0.88 | 0.66 |
| CYS | 0.87 | 0.71 |
| GLN | 0.89 | 0.71 |
| HIS | 0.91 | 0.71 |
| ASN | 0.91 | 0.71 |
| THR | 0.94 | 0.71 |
| SER | 0.95 | 0.71 |
| TYR | 0.95 | 0.78 |
| TRP | 0.96 | 0.78 |
| ALA | 0.97 | 0.78 |
| PHE | 0.98 | 0.78 |
| PRO | 0.98 | 0.78 |
| ILE | 0.99 | 0.78 |
| LEU | 0.99 | 0.78 |
| VAL | 1.00 | 0.78 |

**Supplemental Table 2.** Mitoribosome per-chain refinement results

| Chain ID | iFSC | MolProbity Score | Ramachandran Favored | EMRinger Score |
| --- | --- | --- | --- | --- |
| 0 | 0.649 / 0.646 | 2.85 / 0.84 | 86.17 / 98.94 | 1.74 / 2.60 |
| 1 | 0.653 / 0.647 | 2.48 / 0.77 | 92.00 / 96.00 | 0.86 / 2.75 |
| 2 | 0.711 / 0.688 | 1.81 / 0.84 | 97.56 / 95.12 | 1.68 / 4.15 |
| 3 | 0.730 / 0.698 | 2.11 / 1.06 | 94.62 / 94.62 | 3.11 / 5.09 |
| 4 | 0.707 / 0.653 | 2.29 / 0.50 | 100.00 / 100.00 | 4.32 / 4.40 |
| 5 | 0.696 / 0.688 | 2.76 / 1.31 | 89.40 / 94.02 | 2.80 / 2.68 |
| 6 | 0.593 / 0.563 | 2.49 / 1.39 | 87.54 / 90.73 | 2.54 / 2.20 |
| 7 | 0.648 / 0.637 | 2.27 / 0.98 | 91.47 / 94.19 | 2.22 / 2.14 |
| 8 | 0.378 / 0.359 | 1.67 / 0.80 | 96.36 / 98.18 | 2.03 / 0.68 |
| 9 | 0.626 / 0.591 | 1.91 / 1.61 | 88.57 / 92.38 | 2.25 / 2.19 |
| D | 0.708 / 0.689 | 2.17 / 1.39 | 94.87 / 93.16 | 3.50 / 4.11 |
| E | 0.733 / 0.712 | 2.58 / 1.35 | 90.54 / 94.63 | 2.40 / 3.29 |
| F | 0.718 / 0.698 | 2.50 / 1.02 | 91.94 / 93.55 | 3.71 / 3.44 |
| H | 0.617 / 0.621 | 2.67 / 1.04 | 87.10 / 96.77 | 2.55 / 2.79 |
| I | 0.454 / 0.527 | 2.68 / 0.94 | 92.86 / 97.40 | 2.02 / 1.26 |
| J | 0.334 / 0.420 | 2.73 / 1.40 | 83.33 / 92.75 | -0.12 / 1.41 |
| K | 0.701 / 0.673 | 2.35 / 0.98 | 89.71 / 94.86 | 2.04 / 3.61 |
| L | 0.715 / 0.701 | 2.42 / 1.18 | 88.50 / 94.69 | 0.26 / 1.63 |
| M | 0.711 / 0.687 | 2.55 / 1.47 | 89.82 / 93.33 | 2.04 / 2.43 |
| N | 0.702 / 0.692 | 2.25 / 1.16 | 91.63 / 92.61 | 2.53 / 2.89 |
| O | 0.704 / 0.691 | 2.91 / 1.23 | 88.67 / 96.67 | 1.55 / 3.10 |
| P | 0.675 / 0.659 | 2.72 / 0.99 | 89.15 / 95.35 | 2.22 / 4.80 |
| Q | 0.704 / 0.694 | 2.76 / 0.97 | 89.50 / 96.00 | 2.90 / 3.86 |
| R | 0.704 / 0.687 | 2.31 / 0.95 | 92.75 / 95.65 | 4.19 / 4.51 |
| S | 0.689 / 0.671 | 2.31 / 1.01 | 93.51 / 96.10 | 3.84 / 2.02 |
| T | 0.703 / 0.687 | 2.49 / 0.91 | 92.68 / 96.95 | 2.79 / 2.50 |
| U | 0.699 / 0.697 | 2.59 / 1.06 | 88.07 / 96.33 | 3.20 / 4.32 |
| V | 0.585 / 0.563 | 2.17 / 1.28 | 87.98 / 94.54 | 1.54 / 1.93 |
| W | 0.703 / 0.691 | 2.61 / 0.95 | 97.14 / 96.19 | 3.82 / 6.28 |
| X | 0.672 / 0.656 | 2.56 / 0.50 | 91.29 / 98.34 | 2.12 / 1.74 |
| Y | 0.670 / 0.649 | 2.46 / 1.07 | 90.23 / 95.98 | 2.98 / 3.46 |
| Z | 0.695 / 0.665 | 2.32 / 0.89 | 90.68 / 96.61 | 4.27 / 4.15 |
| a | 0.667 / 0.635 | 2.38 / 1.02 | 94.59 / 97.30 | 1.08 / 5.31 |
| b | 0.690 / 0.658 | 2.46 / 0.91 | 85.62 / 93.84 | 2.42 / 2.48 |
| c | 0.683 / 0.685 | 2.51 / 0.83 | 87.82 / 97.42 | 2.06 / 2.19 |
| d | 0.582 / 0.579 | 1.85 / 1.08 | 89.10 / 96.84 | 2.44 / 3.50 |
| e | 0.163 / 0.279 | 2.74 / 1.75 | 78.03 / 86.36 | -0.03 / 1.44 |
| f | 0.561 / 0.510 | 2.31 / 1.29 | 88.73 / 94.37 | 2.05 / 0.62 |
| g | 0.681 / 0.653 | 2.31 / 0.97 | 91.34 / 96.85 | 3.59 / 3.06 |
| h | 0.526 / 0.517 | 2.43 / 1.06 | 85.42 / 94.79 | 1.76 / 1.83 |
| i | 0.703 / 0.686 | 2.81 / 1.20 | 85.11 / 91.49 | 4.61 / 3.76 |
| j | 0.666 / 0.663 | 2.08 / 0.97 | 95.18 / 97.59 | 3.82 / 3.42 |
| k | 0.367 / 0.520 | 3.43 / 0.74 | 78.05 / 98.78 | 3.36 / 2.83 |
| o | 0.650 / 0.626 | 2.73 / 0.80 | 86.96 / 95.65 | 2.83 / 2.36 |
| p | 0.629 / 0.602 | 1.98 / 1.21 | 92.41 / 92.41 | 2.46 / 1.55 |
| q | 0.546 / 0.551 | 1.82 / 0.66 | 95.24 / 98.41 | 2.40 / 2.40 |
| r | 0.663 / 0.646 | 3.00 / 1.03 | 89.29 / 94.29 | 3.73 / 4.60 |
| s | 0.721 / 0.705 | 2.68 / 1.26 | 89.07 / 93.44 | 2.20 / 1.49 |

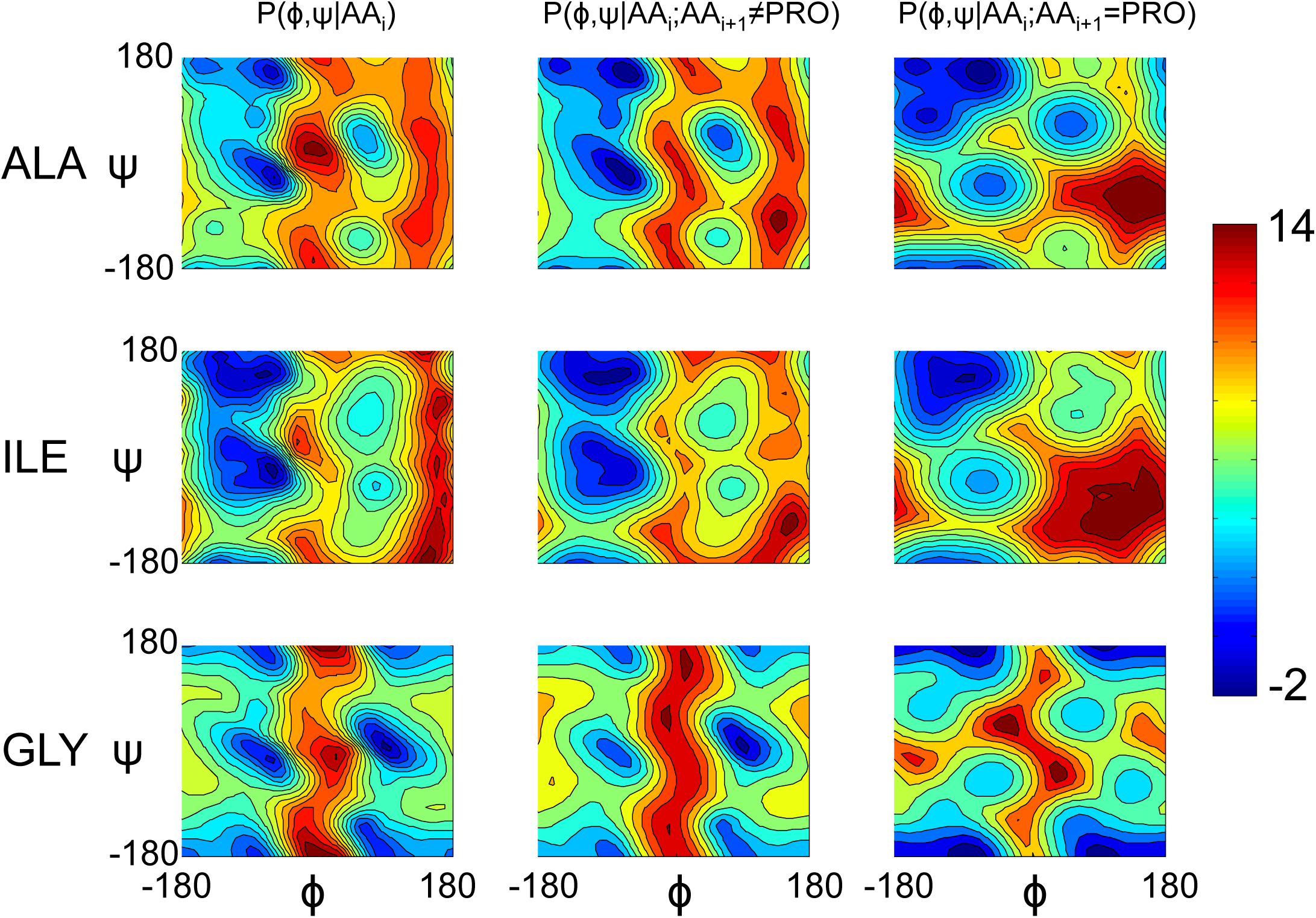

